# *Laocoön*: a tool for high-throughput automated cell counting

**DOI:** 10.1101/751073

**Authors:** Kaitlin Lim, Mikaela Louie, Anne La Torre, Corinne Fairchild, Ian Korf

## Abstract

**Motivation:** There are current programs and plugins that automatically count the number of cells in a given image. However, many of these processes are not entirely automatic, as they require user input to specify a region of interest and are also frequently inaccurate.

**Results:** This project presents *laocoön*, a Python package specifically designed to automatically and efficiently count the number of fluorescently-labelled cells in images. This package not only allows for reliable cell counting, but returns the proportion of cells in each cell cycle relative to all the cells in the DAPI channel, which is currently used for research purposes, but could ultimately be utilized for clinical purposes.

**Availability and Implementation:** This package, its corresponding execution instructions, and further information about the underlying algorithms, are currently available in the GitHub repository https://github.com/edukait/laocoon under the MIT license and can be run on the command terminal of any operating system. Alternatively, *laocoön* is available in the Python Package Index (PyPi), so the user can use the pip command to immediately download the package.

## 1 INTRODUCTION

The cell cycle details the “life cycle” of a eukaryotic cell and is divided into two main parts: interphase and mitosis (Cooper, 2000). Interphase is further divided into a first gap phase (G1), when the cell grows in size synthesizes proteins, the DNA synthesis phase (S), and the second gap phase (G2), which is characterized by cell growth and preparation for mitosis. Mitosis (M) is the process of cell division. Outside of the standard phases in the cell cycle, there is also the G0 phase where the cell remains in a “frozen” state—it is metabolically active, but will not divide unless certain extracellular signals instruct it to do so (Cooper, 2000).

The proper progression of the cell cycle depends on the precise timing of protein synthesis and degradation, as well as the maintenance of the balance between the two (Lecker, Goldberg & Mitch, 2006). The majority of proteins within the cell are degraded by the ubiquitin (Ub)-proteasome pathway (UPP), where enzymes link chains of the polypeptide cofactor Ub on proteins to mark them for degradation. Lecker, Goldberg, and Mitch have determined E3 Ub-protein ligase as the most important enzyme in the process, as it recognizes a specific protein substrate and catalyzes the active transfer of Ub onto said protein. After the enzymes mark the protein, the 26S proteasome then identifies these ubiquitinated proteins, degrading them into small peptides that the cell will eventually reuse (Lecker, Goldberg & Mitch, 2006).

Fluorescent ubiquitination-based cell cycle indicator (FUCCI) cells take advantage of the cell cycle and ubiquitination of certain proteins.. Researchers have attempted to determine how the cell cycle differs depending on the type of cell, environment and interactions. The transition from cell division (M) to G1 is easy to identify by monitoring the morphological changes of the cell. However, the transition from G1 to S is much more challenging to observe in live samples. Prior to the FUCCI labelling technique, researchers observed the transition measured cell cycle by either labelling the cell nuclei with bromodeoxyuridine or synchronizing the cell cycle by pharmacological means (Leonhardt et. al., 2000; Kisielewska, Lu & Whitaker, 2005). Researchers also developed a way to label the cells with fluorescent proteins by fusing them to certain transcriptional proteins, but these markers could not track phase transitions with high contrast (Sakaue-Sawano et. al., 2008).

FUCCI labeling not only provides a quick and relatively cheap means of labelling cells, but can also effectively differentiate between phase transitions. The FUCCI labelling method focuses on two specific substrates of the E3 enzyme: Geminin and Cdt1. Past studies have shown that levels of Geminin and Cdt1 oscillate inversely. While Cdt1 concentration is highest during G1, Geminin concentration is highest during the S, G2, and M phases (Nishitani, Lygerou & Nishimoto, 2004). Researchers were able to fuse red fluorescent protein (RFP) and green fluorescent protein (GFP) to these two substrates to develop fluorescent probes, where cells in the S phase are green, and cells in the G1 phase are red (Sakaue-Sawano et. al., 2008). With this FUCCI labelling technique, researchers can more easily identify whether a cell is in G1 or not.

The training images used in the duration of this project were also labelled with 5-ethynyl-2’-deoxyuridine (EdU). This molecule is a thymidine analogue and can be readily incorporated into cellular DNA during DNA replication, while also allowing good structural preservation of the genetic material (Salic & Mitchison, 2008). During the S phase of the cell cycle, these molecules can interact with fluorescent azides inserted into the cell so they express fluorescence in the image (Salic & Mitchison). EdU labelling combined with FUCCI labelling gives a clearer picture of when and how long the cells are in certain phases of the cell cycle.

The images in the training set are of retinal cells divided into three categories: healthy controls, cells with a let-7 knockout, and cells with let-7 overexpression. The La Torre Lab has chosen to work with retinal cells because their relatively simple architecture and structure allows for easy study of neuronal development and the fate of undifferentiated retinal stem cells, or retinal progenitor cells.

There are six main types of neuronal cells that make up the retina, and all of these cells originate from the same pool of pluripotent retinal progenitor cells (Cepko, 2014; Livesey & Cepko, 2001). This requires careful and precise timing of the proper differentiation of retinal progenitor cells (La Torre, Georgi & Reh, 2013). It has been proposed that retinal stem cells change over time, but the exact underlying mechanisms that explain this phenomenon are unknown. Recent research has shown that microRNAs (miRNAs) play an important role in regulating stem cell differentiation (La Torre, 2013 and Georgi & Reh, 2011). This might in part have to do with the fact that miRNAs also play a crucial role in gene expression by binding to target mRNA to prevent protein production (MacFarlane & Murphy, 2010). Moreover, Fairchild et. al. have also discovered that miRNAs have a role in determining the cell cycle length of retinal progenitor cells (Fairchild et. al., in-press). Subsequently, we have been investigating certain miRNAs for their ability to change cell cycle dynamics in retinal progenitor cells, and let-7 is one of those particular miRNAs under examination (Fairchild et. al.).

Let-7 is one of three types of miRNA that serve as key regulators of the early to late developmental transition in retinal progenitor cells (La Torre, Georgi & Reh, 2013). The levels of Let-7 increase as the retinal progenitor cell fully develops, and it can also partially counteract the dysfunctional development retinal cells that have a knockout of the Dicer complex (Dicer-CKO) (La Torre, Georgi & Reh). Dicer-CKO retinal progenitor cells do not differentiate properly, creating a dysfunctional retina (Damiani et. al., 2008). Interestingly, overexpression of Let-7 together with miR-9 and miR-125b in normal retinas causes accelerated retinal development (La Torre, Georgi & Reh). The objective to see how let-7 expression/knockout influences retinal cell cycle progression inspired the creation of this project—a quicker and easier means of counting cells in many images.

In order to further investigate if cell cycle length can differ between healthy and diseased cells, researchers must take the time to count all the cells in all of the given images. This process is rather long and painstaking and is also subject to human error. The development of *laocoön* to address counting exclusively retinal cells, but the project soon expanded to create a package that can count any and all fluorescently labelled FUCCI and EdU cells. Cell counting is prevalent in many different fields of medicine and research: counting the concentration of red and white blood cells in sick patients, controlling the cell dosage during cell therapy, and much more (Dean, 2005; Simon et. al., 2016). While cell counting in these specific scenarios does not necessarily involve fluorescently labelled FUCCI cells, *laocoön* will hopefully serve as the beginning to a larger-scale project to streamline and automate cell counting.

ImageJ, a Java-based application used primarily for biological image analysis, contains a plug-in for automatic cell counting. However, current researchers often do not use this plug-in because it is often not efficient, and the process of setting up the program and loading the image is tedious. Moreover, developers last updated the plugin in 2010, so some of the algorithms and processes used in the plugin are likely outdated.

This paper aims to show the motivations behind this project, detail the development and testing of *laocoön*, and assess the accuracy of the software.

## 2 TESTING AND DESIGN OF *laocoön*

### 2.1 Training images

The training images were grouped into three categories with 108 images in total. The images were of healthy retinal cells, cells with let-7 knockout, and cells with let-7 overexpression. One snapshot of a slide of cells had four channels, which are effectively four images, and was labeled as one of the following: DAPI, EdU, GFP, or RFP. The DAPI channel included all the cells detected on the slide and served as the reference image for the other three channels. The EdU channel displayed cells that incorporated EdU as they were in S phase. The GFP channel only showed cells that expressed GFP because they were either in the S or G2 phase of the cycle. Lastly, the RFP channel only showed cells that were in the G1 phase of the cell cycle. This combination of EdU labelling with standard FUCCI labelling gave researchers better specificity, as the FUCCI labelling only differentiates the G1 phase from the other phases of the cycle. Figure 1 shows the four different color channels for one slide of cells in which there is let-7 knockout. However, because all of the methods and algorithms can only work with black and white images, all of the images were converted to grayscale before entering the cell counting pipeline.

**Figure 1.**
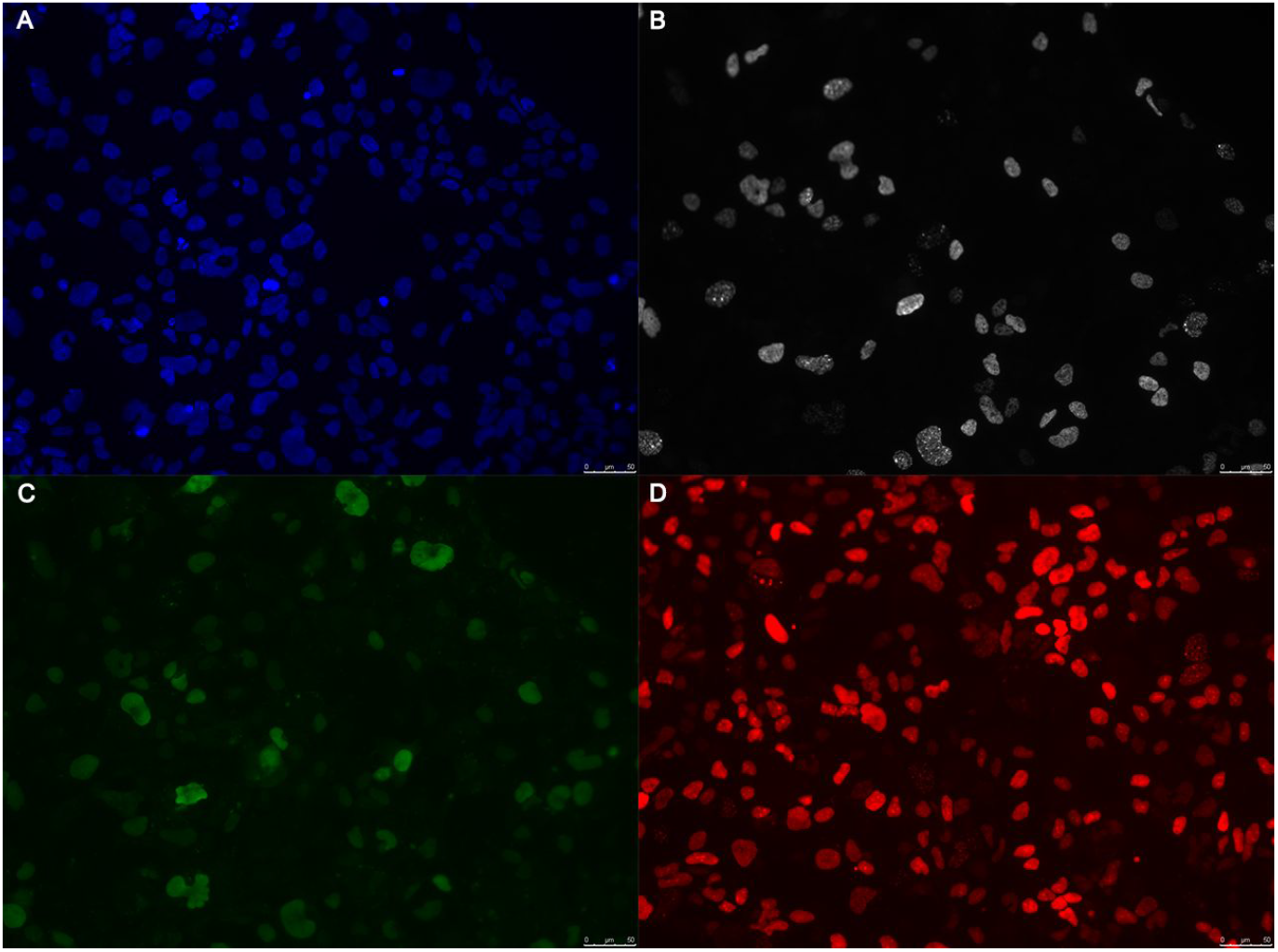
Sample image of the fluorescently labelled cells with let-7 knockout. The figure features the general DAPI channel with all of the cells **(A)**, the EdU-labelled cells **(B)**, the GFP-labelled cells **(C)**, and the RFP-labelled cells **(D)**.

### 2.2 Histogram equalization and connected components

In the process of designing *laocoön*, there were three major algorithms/methods implemented to determine which algorithm would perform best with the training data. The first method implemented histogram equalization and counting the connected components. The former technique preprocessed the image, while the latter actually counted the number of cells in the given image.

Histogram equalization is a technique used on grayscale images to enhance the contrast of the pixels. The histograms represent the distribution of all the pixel values within an image. If *p* is the normalized histogram of *f* with a bin for each possible intensity where the pixel intensities range from 0 to *L* – 1, then

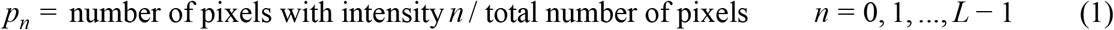

And the histogram equalized image *g* is represented by

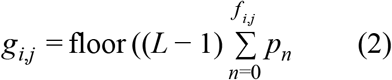

where floor() rounds down to the nearest integer (Gonzalez & Woods, 2008). If a grayscale image is represented as a histogram showing the distribution of pixel values, histogram equalization “stretches out” the histogram, thus enhancing image contrast. Mahotas, the Python package that serves as the main foundation for *laocoön* does not include a built-in function for histogram equalization. As a result, *laocoön* includes its own histogram equalization module.

Following this image preprocessing was counting the cells by finding the number of connected components in the image. This entire methodology is detailed in Rosenfeld and Pfaltz, 1966. The algorithm utilizes the union-find data structure, where all the pixels in the image are looped, or “passed” through, twice. In the first pass, the algorithm goes through all of the pixels. If the computer considers a pixel to be “important” (i.e. not a background pixel), then it looks at the pixels above and to the left of the chosen pixel “p” to determine one of two cases. First, if the pixel above and/or left of it are not background pixels, the algorithm labels it with the same “name” as the other non-background pixels around it. Second, if the pixel is surrounded by background pixels, the algorithm creates a new label that is unique from all the other preexisting labels. In the situation where two different labels are labeling the same component, then, using the union-find structure, the computer sets one of the labeled groups as the “child” of the other labeled group, thus declaring that they are the same group. The algorithm continues to do this until all of the pixels have been passed through. Then, in the second pass through all the pixels, the computer labels all of the “children” of the root label the same label as the parent’s label. As a result, the total number of groups at the end of the process determines the total number of cells in the image.

### 2.3 Gaussian filter and regmax

The second method employed a different form of preprocessing the images and counting the cells. Rather than using histogram equalization, a Gaussian filter was added in order to reduce image noise and detail. As a result, the computer had a smaller probability of confusing a non-cellular structure as a cell. In two dimensions, the Gaussian filter can be represented mathematically as

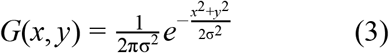

where σ is the standard deviation of the image pixels (Shapiro & Stockman, 2000). The Gaussian filter acted as a convolution on the image, changing the pixel values and subsequently “smoothing” the image (Davies, 2012). Mahotas has a built-in Gaussian filter function, where the user can input a specialized sigma, or standard deviation, that the Gaussian filter uses (Coelho, 2013). This sigma value is unique for each image channel and was determined using qualitative analysis of the resulting image.

After the preprocessing step, the regmax function, also in the Mahotas library, located the perceived centers of the cells in the images. This regmax function finds the regional maxima in the image. In the case of two-dimensional grayscale images, the regional maxima are the connected components of pixels with the same intensity value, which are then surrounded by pixels with lower intensity values (Dinstein & Fong-Lochovsky, 1988). The regmax function returns a matrix of binary values—the regional maxima are marked with 1’s, while everything else is marked with 0’s. Because the cells were of higher pixel intensity values compared to the backdrop, the Gaussian filter served to, in ways, enhance this contrast and thus make the regmax function much more accurate. Rather than identifying the outlines of the cells with the connected components method, which can confuse two overlapping cells as one massive cell, this regmax method identified the centers of these cells and was much more efficient in identifying overlapping cells. As a result, *laocoön* only features regmax counting and not connected component counting because it is the more accurate of the two processes.

### 2.4 Epsilon value quality control

The third and final method involved was an additional quality control component. Neither of the two previously described pipelines had any form of quality control to ensure that the cells in the EdU, GFP, and RFP channels actually existed in the DAPI channel so that the computer was not counting pseudo-cells. More specifically, the regmax algorithm could determine different “centers” of one cell in two different channels. For instance, the cell “center” in the DAPI channel might be right in the middle of the cell, but the perceived “center” in the EdU channel of the very same cell can be on the right end of the cell. The epsilon value addressed this problem. This value served as a form of “padding,” where if the perceived center of the DAPI channel was within that determined radius of the other channel’s “center,” then the cell was not a pseudo-cell and could be counted.

Each channel had different pixel intensity values, and so the maximum distance value is different for each channel, although all channels use the original DAPI channel as the reference image. Because of this, the epsilon value was also determined using qualitative analysis, like for the Gaussian filter’s sigma value.

## 3 IMPLEMENTATION

The *laocoön* package features the Gaussian filter and histogram equalization preprocessing, regmax cell counting, and epsilon quality control. Execution of the main default script creates four cell counting pipelines, all of which implement Gaussian filter preprocessing and regmax counting:

1. Histogram equalization preprocessing; epsilon value quality control
2. Only histogram equalization
3. Only epsilon value quality control
4. Neither

This allows for a diversity of methods of cell counting and a larger arsenal of cell counting tools for the user, as only one pipeline does not guarantee accurate counting for all types of fluorescently labelled cell images.

In order for the script to run properly, there are a few guidelines the input images must follow. The user must download a few Python packages to ensure that the *laocoön* package is functional. All of the images for each labelled collection of cells—DAPI cells, EdU-labelled cells, GFP-labelled cells, and RFP-labelled cells—must also have the same group name and specify which channel the image is in the file name. The GitHub repository also contains instructions on the execution of *laocoön*, as well as images of sample command line inputs in the repository README file.

## 4 DISCUSSION

### 4.1 Results with training data

The sensitivity and specificity metrics of the four pipelines could not be discerned because the training data did not have labeled pixels. Thus, average percent of experimental error was the only means of assessing the accuracy of the code. Figure 4 shows the average percent error across the four different pipelines for the channel counts, as well as the ratios.

**Figure 2.**
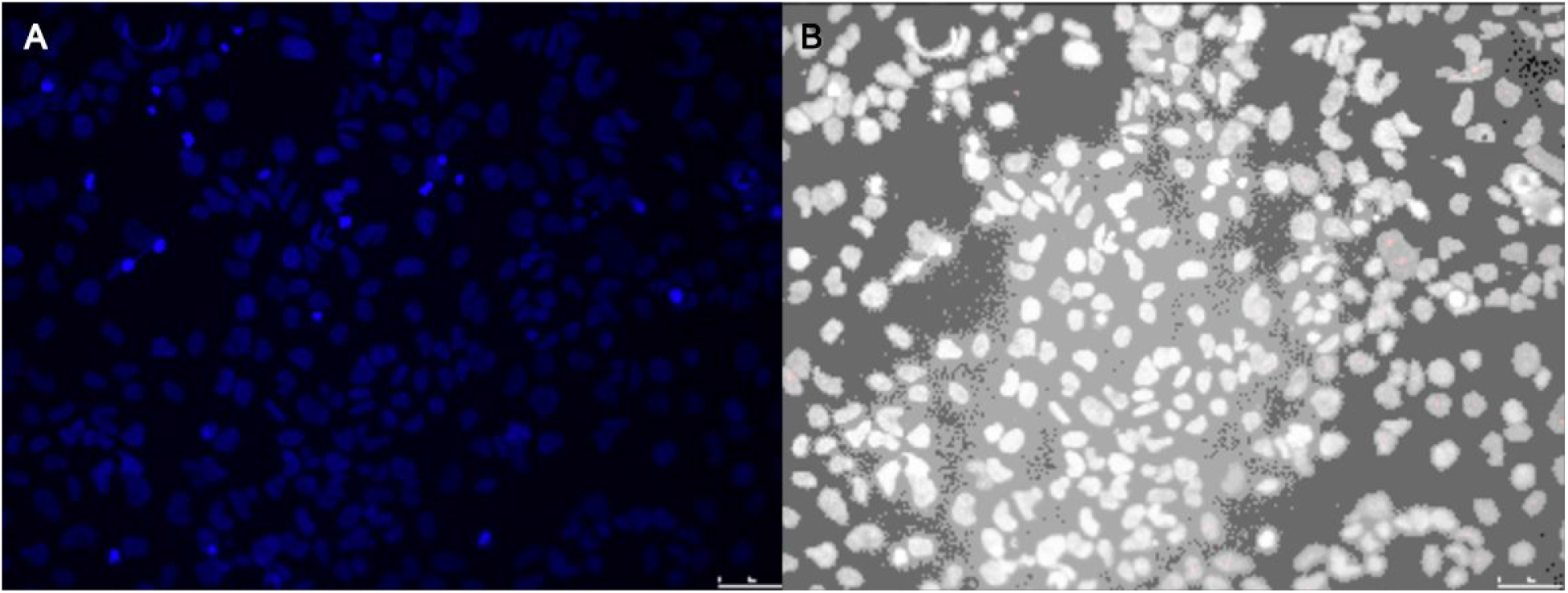
Example of histogram equalization. The figure shows the original image **(A)**, a slide of retinal cells in the DAPI channel, compared to the image after histogram equalization **(B)**.

**Figure 3.**
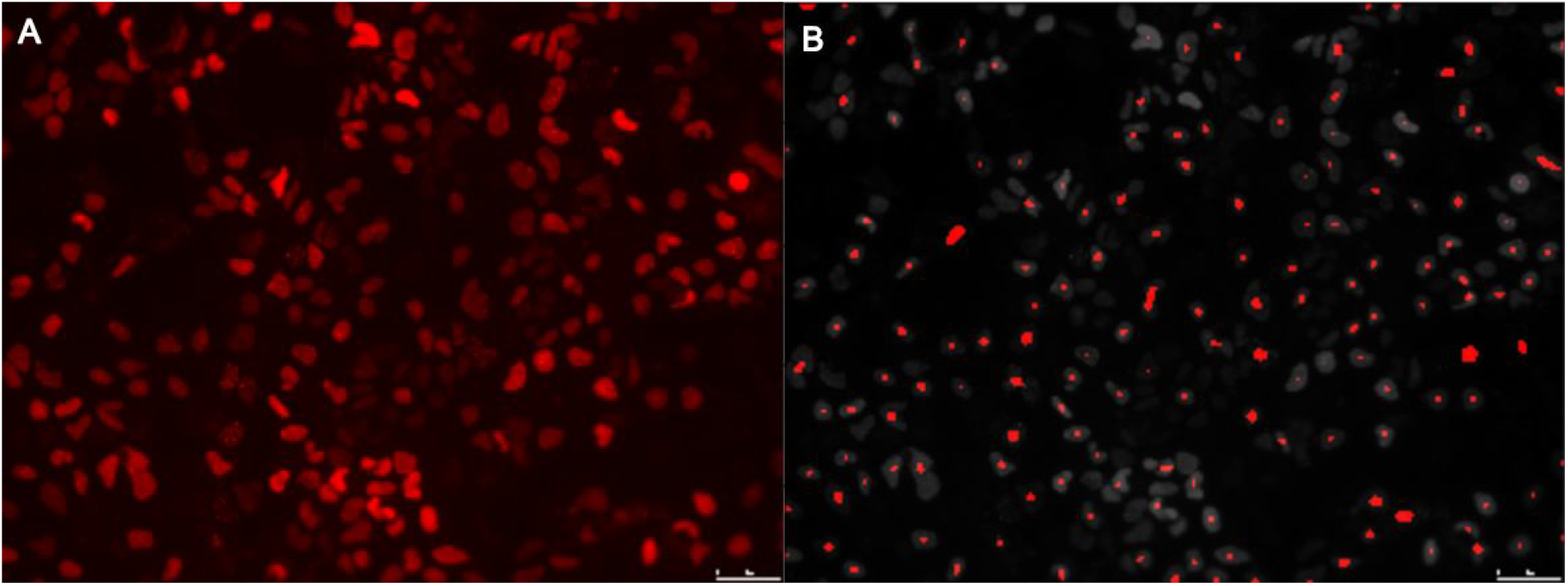
Example of how the regmax algorithm detects the “centers” of cells. The figure shows the original image **(A)**, a slide of let-7 knockout retinal cells in the RFP channel, compared to the regmax-processed image **(B).**

**Figure 4.**
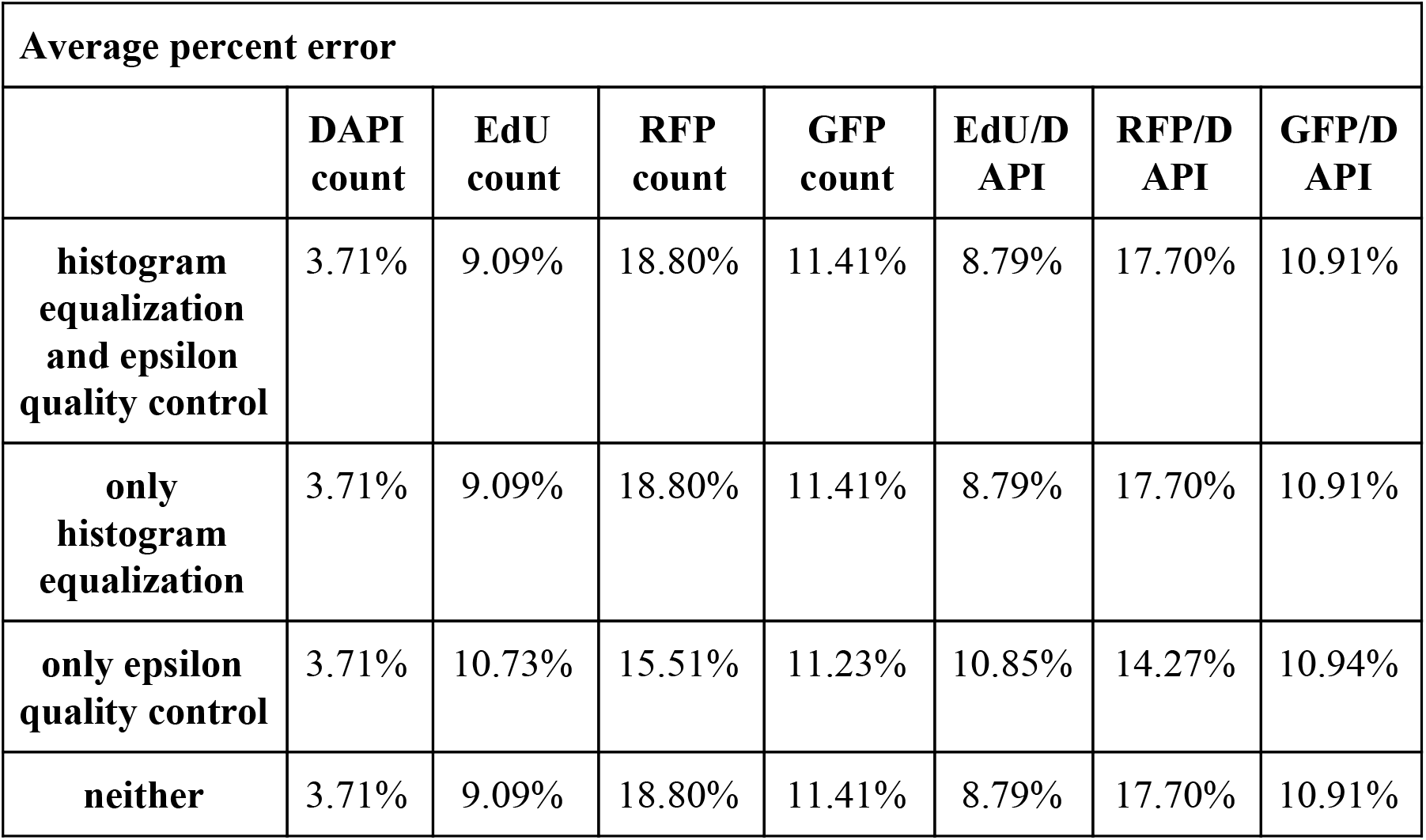
Table displaying the average percent error across the four implemented methods.

The average experimental error for the DAPI channel does not seem to change depending on the pipeline used. This implies that only a Gaussian filter is the only necessary measure taken, and there needs to be no extra preprocessing or quality control.

What was especially surprising was that the average experimental error for the DAPI channel—the image with the largest amount of cells—was the smallest, while the RFP and GFP channels—the images with the most well-defined cells—had the largest average experimental error. This is likely to do with the fact that the cells in the RFP and GFP channels exhibit a wider variation of pixel brightness when compared to the cells in the DAPI channel. In Figure 1, the DAPI channel has a more uniform distribution of pixel brightness across the cells with the occasional bright specks. However, in the RFP and GFP channels, there is a wider variation of pixel brightness among the cells. Some cells are extremely bright, while others seem like faded blots in comparison. This likely affects the preprocessing of RFP and GFP channel images. Because histogram equalization increases the contrast within the image, that might make the already dim cells seem even dimmer, so the regmax function subsequently does not identify them as cells.

Generally speaking, the differences in the average experimental error seems more indicative of greater differences between the channels rather than the four pipelines. As a result, this might require further development on preprocessing the images—especially the images in the RFP and GFP channels, as they have the highest average experimental error.

### 4.2 Future directions

One future direction is to improve the software’s quality control and overall accuracy—especially in RFP channel counting and GFP channel counting—possibly with a more complex, advanced structure such as convolutional neural networks. It might also be worth investigating how other forms of image preprocessing can enhance the cell counting performance on RFP and GFP channel images with, for instance, decreasing the contrast rather than enhancing it.

Another future direction, albeit a far more ambitious one, is to create an entirely new library that implements methods and classes without the need for external packages. This would allow for more independence within the way the algorithms are written and the data structures are used in the software. This added flexibility can also reduce the memory used during and time complexity of running programs. Similarly, *laocoön* can become a package for more than just fluorescently labelled cell counting and expand into other types of cell images, and the creation of an entirely new library can aid in this initiative.

Creating a new means of automatic cell counting in the form of a Python package will increase the accessibility and ubiquity of these types of tools. *Laocoön* hopes to introduce an updated method of cell counting that can handle batches of images at a given time while also being efficient. This software package aims to make cell counting an easier process and subsequently aims to expedite research and clinical studies involving these types of images.

## ACKNOWLEDGEMENTS

I would like to thank Ian Korf for providing me the opportunity to work with him over the summer at the University of California, Davis, as well as giving me advice on what I can do to further improve my code. I would also like to thank Mikaela Louie, who first introduced me to this project, and Anne La Torre, who gave me access to her cell images for training data.

